# The Easter Egg Weevil (*Pachyrhynchus*) genome reveals synteny in Coleoptera across 200 million years of evolution

**DOI:** 10.1101/2020.12.18.422986

**Authors:** Matthew H. Van Dam, Analyn Anzano Cabras, James B. Henderson, Cynthia Pérez Estrada, Arina D. Omer, Olga Dudchenko, Erez Lieberman Aiden, Athena W. Lam

## Abstract

Patterns of genomic architecture across insects remain largely undocumented or decoupled from a broader phylogenetic context. For instance, it is unknown whether translocation rates differ between insect orders? We address broad scale patterns of genome architecture across Insecta by examining synteny in a phylogenetic framework from open source insect genomes. To accomplish this, we add a chromosome level genome to a crucial lineage, Coleoptera. Our assembly of the *Pachyrhynchus sulphureomaculatus* genome is the first chromosome scale genome for the hyperdiverse Phytophaga lineage and currently the largest insect genome assembled to this scale. The genome is significantly larger than those of other weevils, and this increase in size is caused by repetitive elements. Our results also indicate that, among beetles, there are instances of long-lasting (>200 Ma) localization of genes to a particular chromosome with few translocation events. While some chromosomes have a paucity of translocations, intra-chromosomal synteny was almost absent, with gene order thoroughly shuffled along a chromosome. To place our findings in an evolutionary context, we compared syntenic patterns across Insecta. We find that synteny largely scales with clade age, with younger clades, such as Lepidoptera, having especially high synteny. However, we do find subtle differences in the maintenance of synteny and its rate of decay among the insect orders.

## INTRODUCTION

Beetles represent one of the most diverse groups of metazoans, with ∼400,000 described species (Hammond 1992), and estimates of total diversity up to 0.9–2.1 million species (Stork et al. 2015). Among beetles, weevils (Coleoptera: Curculionidae) are one of the most diverse insect groups (>60,000 species (Oberprieler et al. 2007)), encompassing a huge range of life history strategies and occupying every conceivable niche in a terrestrial ecosystem. With morphological forms specialized to ecological habits, such as feeding on fungi, seeds, pollen, wood, roots, and even kangaroo dung, weevils make an excellent system in which to study the evolution of different ecomorphologies (Zimmerman 1994, Oberprieler et al. 2007). Weevils belong to the group Phytophaga whose members comprise lineages that specialize on and have co-diversified with many plant lineages (McKenna et al. 2009, Seppey et al. 2019). Given their vast diversity and economic importance as pollinators and crop pests, knowing more about the genomic architecture of beetles should be of broad applicability. However, to date, there are only four available genomes resolved to chromosome level for Coleoptera and none for weevils or the hyperdiverse beetle lineage Phytophaga (Van Belleghem et al. 2018, Fallon et al 2018, Zhang et al. 2020, Herndon et al. 2020). Here we present the first genome resolved to chromosome level for the Phytophaga beetle lineage *Pachyrhynchus sulphureomaculatus* Schultze, 1922.

Recent advances in genome assembly techniques, such as in situ high throughput conformation capture technology (Hi-C) (Lieberman-Aiden et al. 2009), have substantially enhanced our knowledge of genome architecture (Rao et al. 2014, Dudchenko et al. 2017, Eagen et al. 2017). Increasing the accuracy and contiguity of genome assemblies has also been aided by using long-read sequencing technology in combination with in situ Hi-C (Ghurye et al. 2017, Matthews et al. 2018, Song et al. 2018, Kingan et al. 2019, Scheffer et al. 2020). These innovations have allowed researchers to not only reconstruct genomes to chromosome scale but also to do so relatively quickly and cheaply (Dudchenko et al. 2018). In addition, in situ Hi-C technology has shown that the 3D conformation of genomes is not random and that this conformation can influence gene expression and linkage (Sanborn et al. 2015). The result of these new sequencing techniques has increased the number of high quality genomes for non-model insect species, including beetles (Matthews et al. 2018, Hill et al. 2019, Lu et al. 2019, Liu et al. 2019, Biello et al. 2020, Herndon et al. 2020, Zhang et al. 2020). Because in situ Hi-C orders scaffolds and corrects misjoins, we can study synteny between organisms with more confidence (Dudchenko et al. 2017, Ghurye et al. 2019).

With the influx of new chromosome-level genomes, we can now begin to explore patterns of genome architecture within and between major insect lineages. For example, in Lepidoptera (butterflies), genome architecture has been characterized as relatively stable with few (6%) orthologous loci being translocated (Ahola et al. 2014, Davey et al. 2016, Hill et al. 2019, Wan et al. 2019). Holocentric chromosomes observed throughout Lepidoptera are implicated in reducing hybridization limitations, (Marec et al. 2001, Lukhtanov et al. 2018, Edelman et al. 2019, Hill et al. 2019) suggesting that genome architecture plays a significant role in their biology. In *Heliconius* butterflies, the inversion and rearrangements that do occur do not seem to hinder hybridization (Edelman et al. 2019). In contrast to Lepidoptera, *Drosophila* species have many more translocations and rearrangements (Renschler et al. 2019). In beetles, however, even a basic understanding of genomic architecture remains undocumented. The basic blueprints as revealed by in situ Hi-C maps of how a genome is organized (e.g. – with a Rabl-like conformation (Rabl 1885, Csink and Henikoff 1998), holocentric chromosomes, chromosome domain territories, compartments, and topological associated domain loops) remain non-existent and therefore unplaced in a phylogenetic context. A general synthesis across insects linking these genomic architectural patterns to their function and potential influence on speciation remains incomplete. For example, do different insect orders have distinct rates of genomic rearrangements (the breakage of synteny between genes), or are the patterns we observe merely due to clade age? Are there aspects of a lineages’ genomic architecture that contribute to their observed syntenic patterns? Here we address these questions and provide a new chromosome-level genome for Coleoptera.

## RESULTS

### Sequencing and assembly results

From our PacBio library we sequenced a total of 87,452,300,317 base pairs (bp) with an N50 read length of 31,404 bp (see Supplemental Information Table 1 for full report). From our in situ Hi-C library (we refer to the in situ Hi-C library or reads as “Hi-C” throughout), we sequenced a total of 228,169,567 paired reads after cleaning. Only 2.53% of our Hi-C reads were unmapped, and we had a total of 80,652,881 Hi-C contacts. For a list of the intra-/inter-chromosomal contacts and long/short range Hi-C contacts, see Table 1.

**Table 1.** Summary of Hi-C reads mapped.

|  |
| --- |
| Inter-chromosomal: 56,711,177 (24.85% / 38.63%) |
| Intra-chromosomal: 23,941,704 (10.49% / 16.31%) |
| Short Range (<20Kb): 18,227,329 (7.99% / 12.42%) |
| Long Range (>20Kb): 5,714,353 (2.50% / 3.89%) |

Our initial assembly after 3X polishing in RACON (Vaser 2017) consisted of 18,240 scaffolds and was 2,982,578,979 bp in total length. After removing duplicate haplotigs with *Purge Haplotigs* (Roach et al. 2018), 9,751 scaffolds and 2,052,097,903 bp remained (Fig. 2). Our initial Hi-C assembly resulted in 4,111 and 2,057,226,403 bp total. The size increase is due to 500 bp insertions of Ns (the 3D-DNA default), between scaffolds merged into super-scaffolds. Running *Pilon* (v. 1.23) (Walker et al. 2014) in “--fix bases’’ mode and removal of mitochondrial and contaminant scaffolds (virus or bacteria) resulted in 4,093 scaffolds and 2,051,389,195 bp. A bubble plot of scaffolds by taxon category, and a table of the chromosomes scaffold N50s are shown in Fig. 3 and Table 2, respectively. The identity of other scaffolds not included in the main chromosomes are ambiguous (14 potential viruses and 31 potential bacteria). We retained these but did remove any with bacteria or virus as their best blast score those previously mentioned. The full summary statistics of our final assembly are shown in Table 2. From the different versions of BUSCO (Felipe et al. 2015) Insecta gene sets (1658 BUSCOs version 2, 1367 version 4 and 5-beta), the percentage of complete genes varied (90.8% V2, 87.6% V4 and 91.1% V5 (Fig. 4)), indicating a relatively complete assembly. Compared to other chromosome-level beetle genomes, we found a comparable number of complete BUSCO genes. However, the results vary somewhat depending on which version of BUSCO and which genes were used (Fig. 4). We found a relatively low duplication rate compared to that found in two other beetle (*Photinus* firefly and *Propylea* ladybeetle) genomes that used primarily long-read and Hi-C sequencing in their assembly.

**Table 2.** Summary statistics for final assembly.

|  |  |
| --- | --- |
| Number of scaffolds | 4,093 |
| Total size of scaffolds | 2,051,389,195 |
| Number of contigs | 14,365 |
| Number of contigs in scaffolds | 10,283 |
| Number of contigs not in scaffolds | 4,082 |
| Mean scaffold size | 501,195 |
| Median scaffold size | 8,175 |
| N50 scaffold length | 215,921,627 |
| scaffold %AT | 33.11 |
| scaffold %CG | 16.87 |
| scaffold %N | 0.05 |
| **% bp of assembly in chromosomes** | **97.52** |

**Table 3.** Summary statistics for final assembly by chromosome.

| Chromosome | length bp | # of contigs | number of N's (runs of 100) | percent N's in chromo. | N50 | N50 reached in # of contigs |
| --- | --- | --- | --- | --- | --- | --- |
| Chr_1 | 263,832,947 | 1388 | 138,700 | 0.05% | 287319 | 280 |
| Chr_2 | 253,284,860 | 1222 | 122,100 | 0.04% | 326447 | 246 |
| Chr_3 | 137,890,936 | 683 | 68,200 | 0.04% | 315991 | 127 |
| Chr_4 | 223,502,247 | 1131 | 113,000 | 0.05% | 297265 | 217 |
| Chr_5 | 69,931,891 | 236 | 23,500 | 0.03% | 452330 | 50 |
| Chr_6 | 125,299,487 | 655 | 65,400 | 0.05% | 304080 | 131 |
| Chr_7 | 132,078,125 | 624 | 62,300 | 0.04% | 368684 | 112 |
| Chr_8 | 173,282,956 | 836 | 83,500 | 0.04% | 335626 | 165 |
| Chr_9 | 215,921,627 | 1221 | 122,000 | 0.05% | 280522 | 237 |
| Chr_10 | 186,927,849 | 988 | 98,700 | 0.05% | 290475 | 197 |
| Chr_11 | 218,628,933 | 1074 | 107,300 | 0.04% | 316887 | 219 |

**Figure 1.**
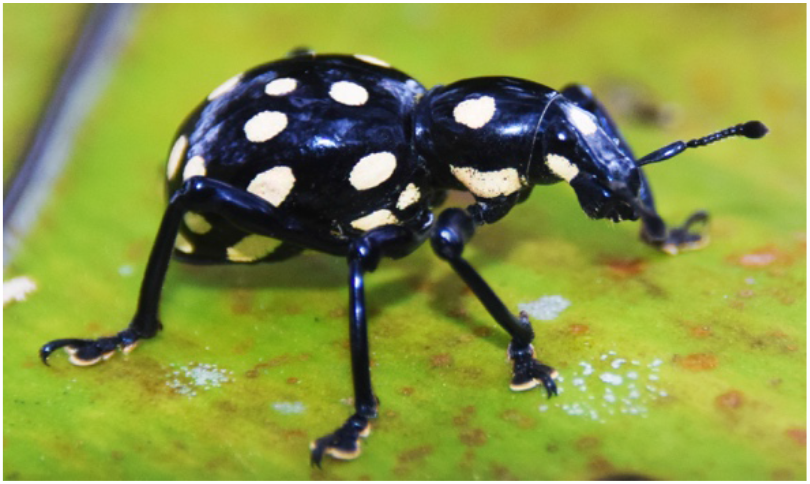
*Pachyrhynchus sulphureomaculatus*, lateral habitus. (Photo by A. Cabras)

**Figure 2.**
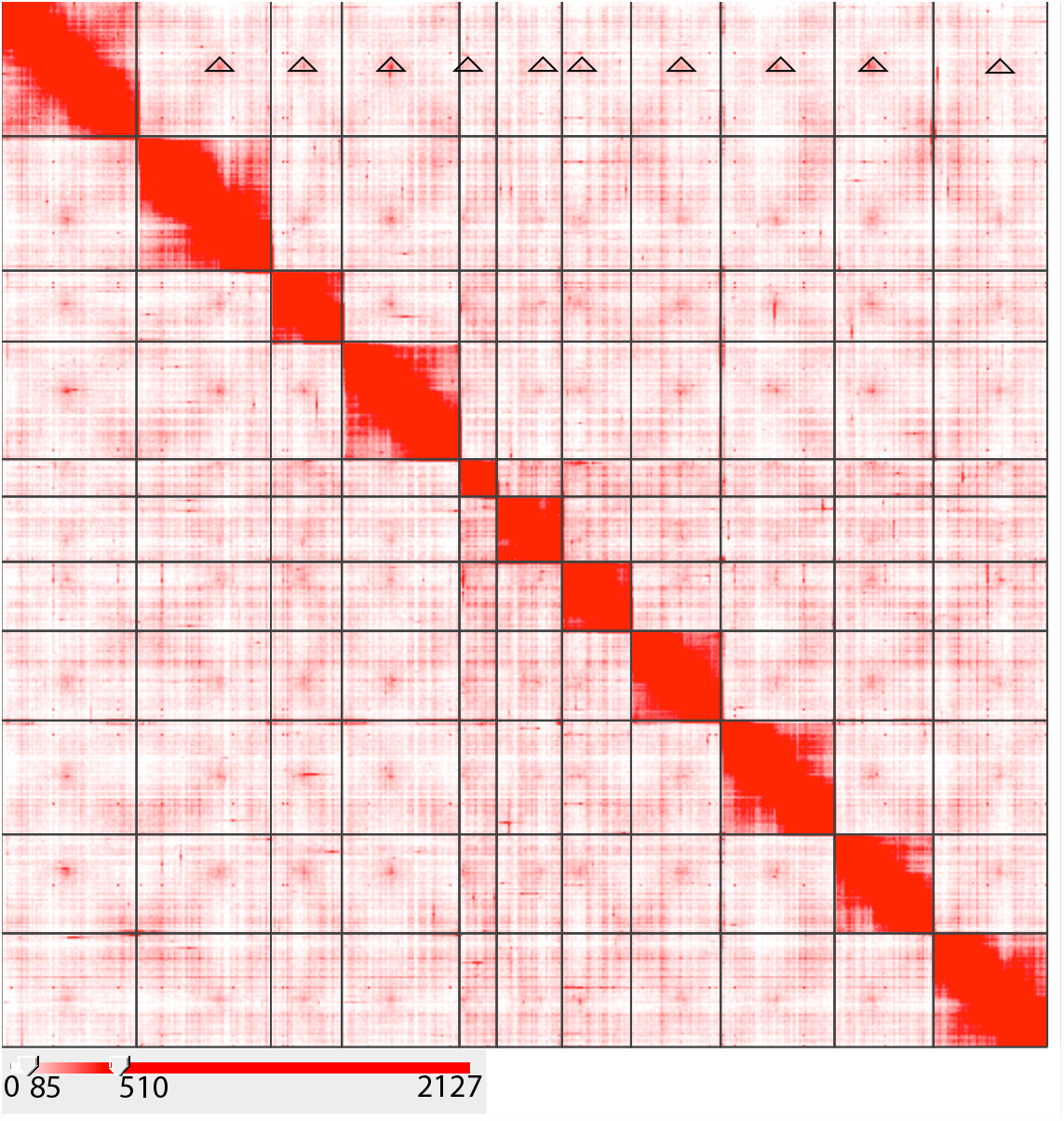
Hi-C contact map heatmap of *Pachyrhynchus sulphureomaculatus* Schultze, 1922. Eleven chromosome boundaries are indicated by black lines. Heatmap scale lower left, range in counts of mapped Hi-C reads per megabase squared. Rabl-like pattern highlighted along chromosome 1, top row, open triangles indicate contact between centromere regions. X-like pattern between adjacent off diagonal regions indicative of contact between distal portions of chromosomes.

**Figure 3.**
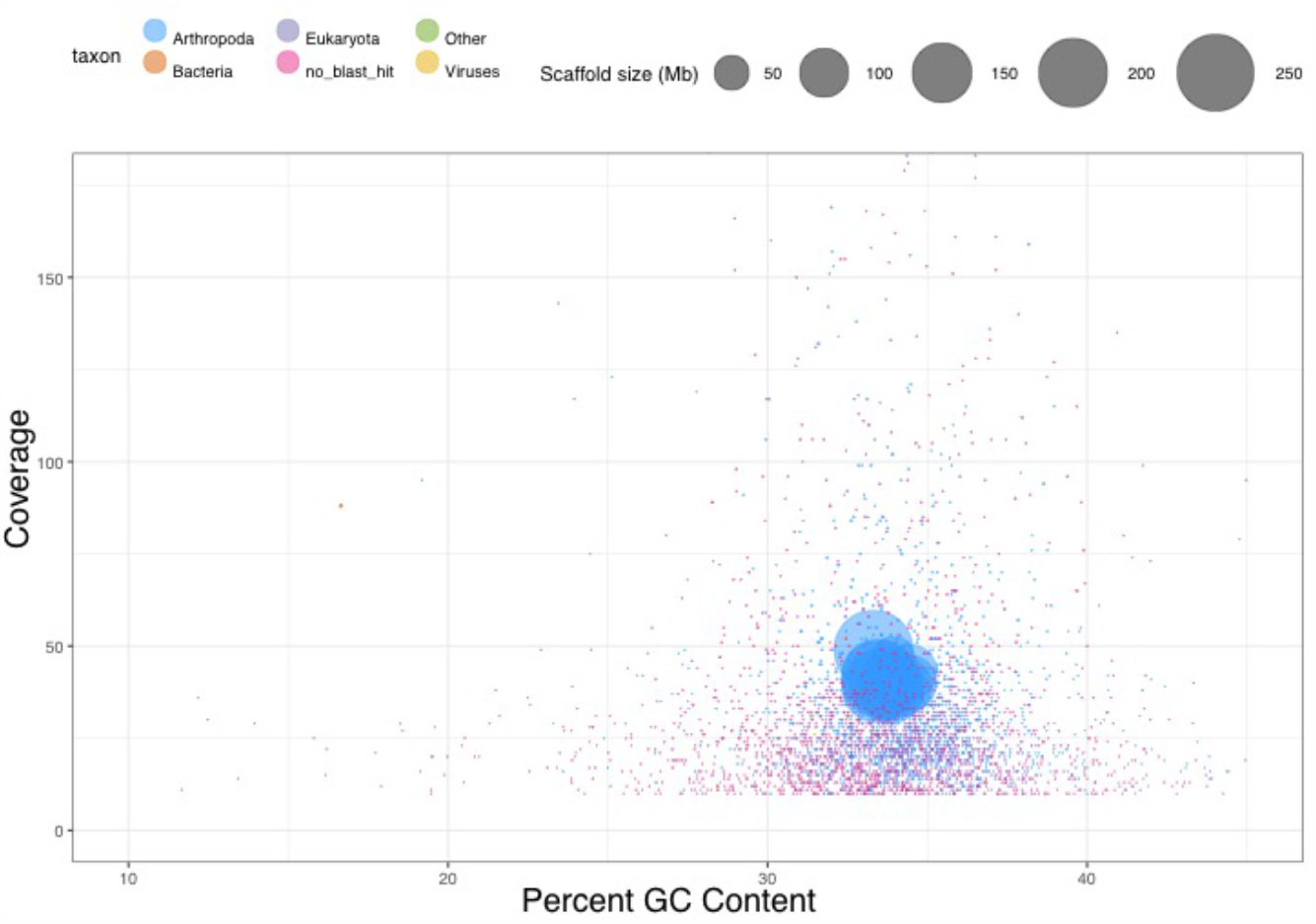
*P. sulphureomaculatus* scaffold bubble plot of coverage versus GC content. Scaffolds included are from the unfiltered assembly. Taxonomic annotation provided via *blastn* alignment to the NCBI nt database.

**Figure 4.**
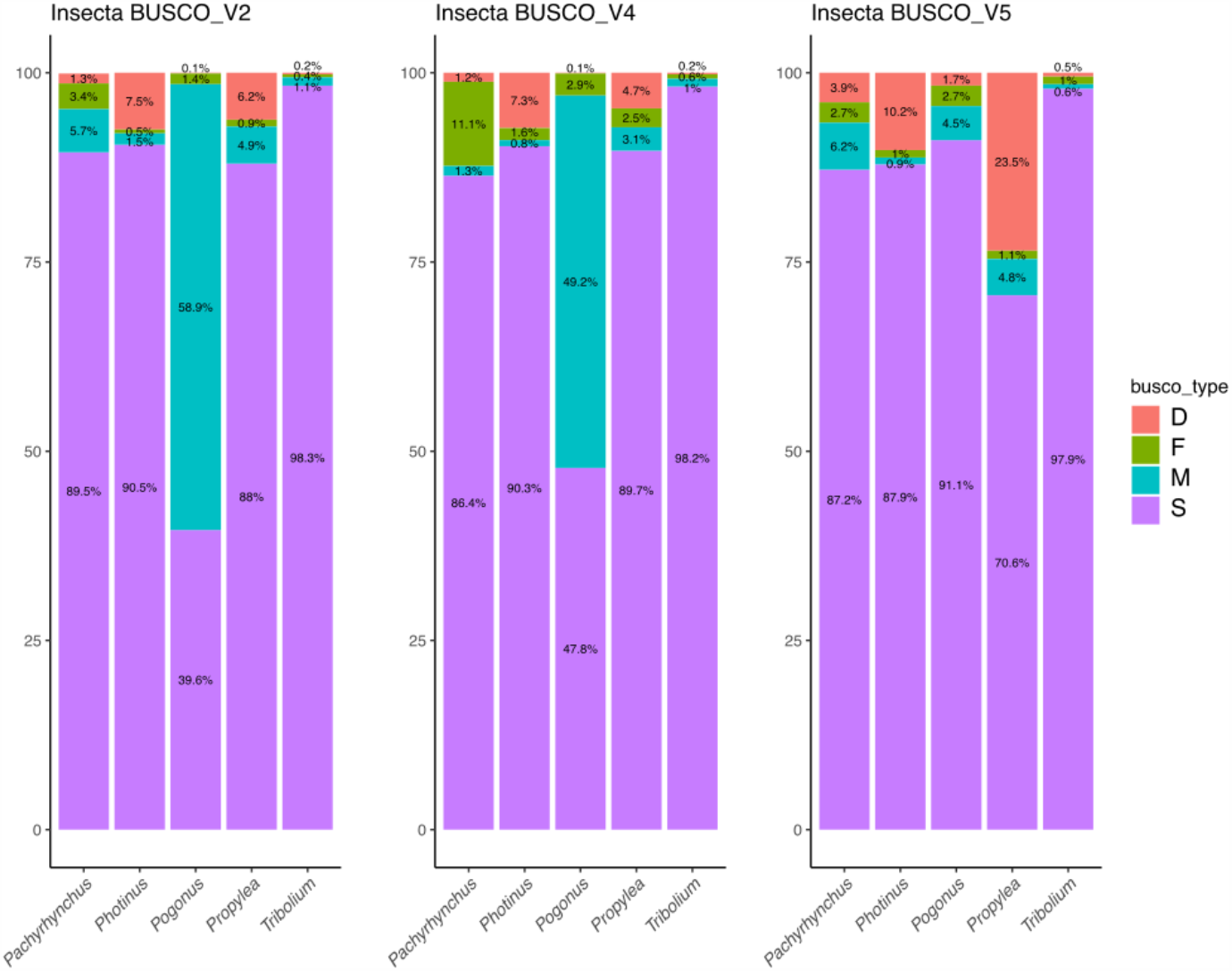
Stacked bar plot of Insecta BUSCO gene sets by category for chromosome-level beetle genomes. Y-axis is the percent of BUSCO genes, X-axis labels are the genus names. The abbreviations in the legend are: D=duplicated, F=fragmented, M=missing and S=single.

### Repeat content analyses

At 2.05 Gbp, the *Pachyrhynchus sulphureomaculatus* genome is roughly 1.8 times as large as the next largest weevil (Curculionoidea) genome published to date, the 1.11 Gbp *Listronotus bonariensis*, the Argentine Stem Weevil (Harrop et al. 2020), and 2.6 times the next largest, the 782 Mbp Red Palm Weevil, *Rhynchophorus ferrugineus* (Hazzouri et al. 2020) genome. To help explain the size difference, we categorized the repeat content of *P. sulphureomaculatus*. The repeat content analyses from *RepeatMasker* shows that the genome of *P. sulphureomaculatus* consists of more than three quarters (76.36%) repetitive DNA, similar to the repeat percentage of *Listronotus*, which is the closest relative to *Pachyrhynchus*. Compared to other weevil genomes (Fig. 5), *P. sulphureomaculatus* has roughly the same percentage of non-repetitive DNA as *Listronotus* and *Sitophilus*. However, the genomes of the two bark beetles of the subfamily Scolytinae (*Dendroctonus* and *Hypothenemus*), are ∼1/12 the size of *P. sulphureomaculatus* and consist of only ∼17% repetitive content. The *P. sulphureomaculatus* genome consisted of 73.1% interspersed repeats, with SINEs being 0.1%, LINEs 20.8%, LTR elements 2.6%, DNA elements 33% and unclassified repeats 16.6%. A sliding window analysis suggests that repetitive content tends to be found in a higher percentage towards the ends of the chromosomes in *P. sulphureomaculatus*, except in chromosome 5 (Fig. 6).

**Figure 5.**
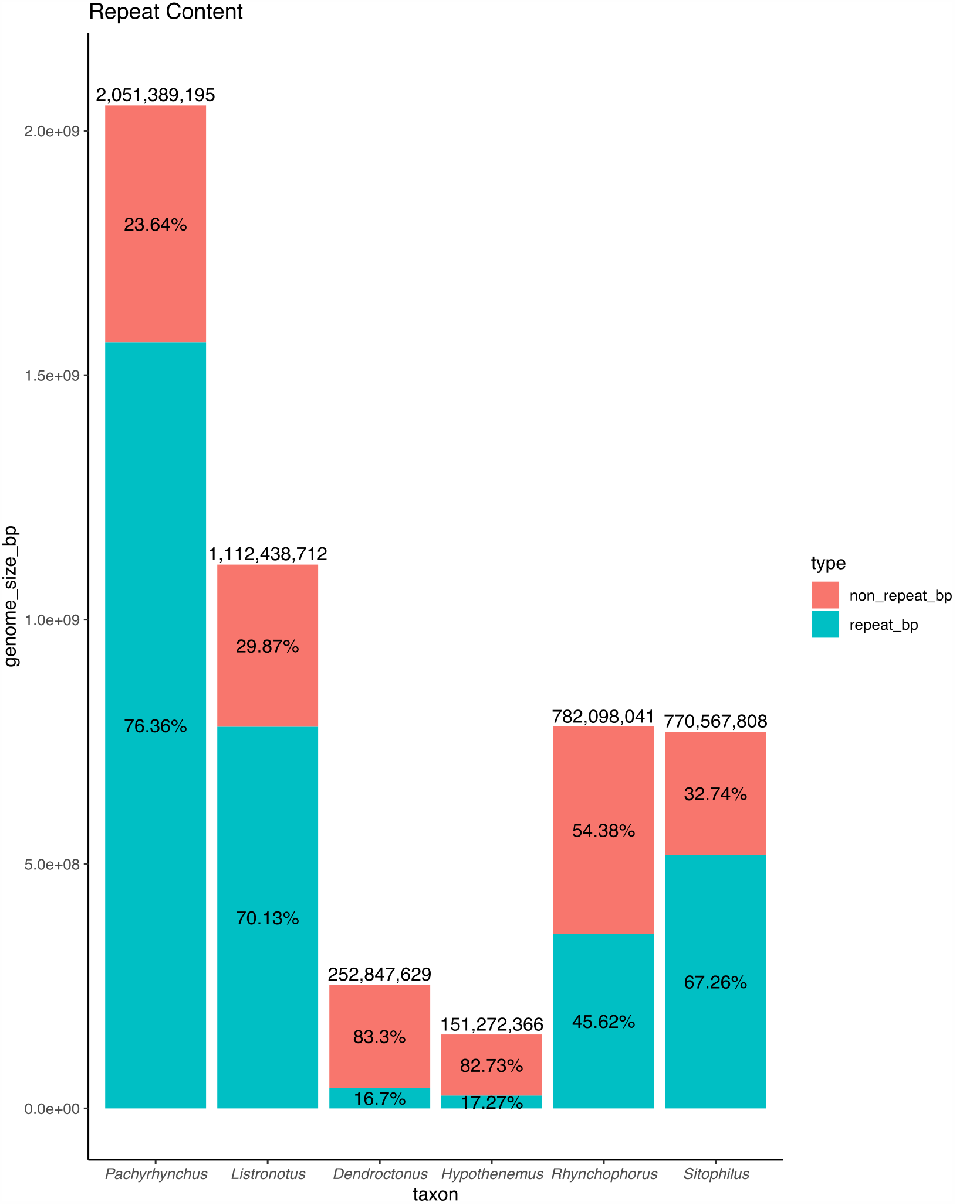
Histogram of repeat content for weevil genomes.

**Figure 6.**
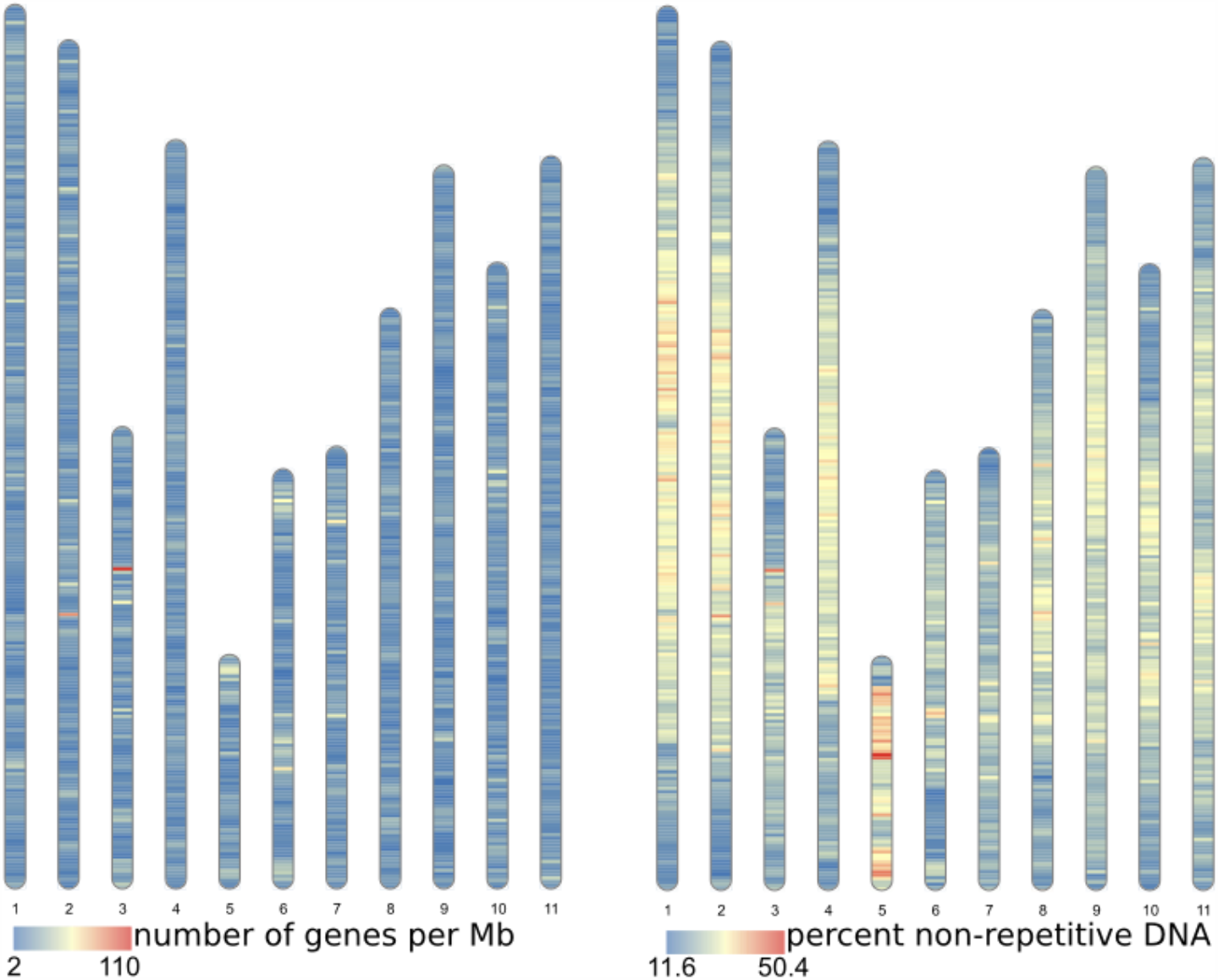
Heat map of gene density and non-repetitive DNA per 1 Mb sliding window. The 11 chromosomes are in the same order as in the Hi-C heat map (Fig. 1) and fasta file.

### Genome annotation

After removing low quality reads from our transcriptome library, a total of 20,551,938 paired reads remained. Our initial 3 transcriptome assemblies, *Trinity* de novo, *Trinity* genome guided assembly and *rnaSPAdes*, resulted in fairly similar assemblies, with each having a high number (∼90%) of the BUSCO v.2 Arthropoda genes (see Supplemental Information Table 2 for details).

As the nuclei of cells between different species generally do not interact (except for viruses), and because Hi-C mapping will remove any non-*Pachyrhynchus* DNA from the chromosomes, we only annotated genes found within the 11 chromosomes comprising 2,000,581,858 bp. The *EVidenceModeler* analysis found that the *P. sulphureomaculatus* contained, as percentages of length, 26.00% genes, 1.46% exons and 24.54% introns, with 30,175 genes, totaling 520,120,665 bp with a mean length of 17,236.81. There are a total of 10,009 single exon genes (33.17% of genes), a total of 120,454 exons with a mean of 3.99 exons per gene, a length of 242.1 bp and per gene length of 966.41bp. There are a total of 90,279 introns with a mean of 2.99 per gene, a length of 5,438.24 bp and a per gene length of 16,270.4 bp. The distribution of gene, exon and intron sizes can be found in the Supplemental Information file “Size_of_Genes_Exons_Introns”. The gff and tRNA annotations are also in the Supplemental Information. Chromosome gene distribution is relatively even, with only a few regions enriched with genes (Fig. 6).

### Synteny across coleopteran chromosome-level genomes

We found BUSCO v.2 loci (1658 Insecta gene set) had a low level of translocations between chromosomes (Fig. 7). Our UCE set resulted in 295 loci among taxa and recovered an identical topology and similar dates as in McKenna et al. (2019). We also found a similar synteny pattern between BUSCO genes and those from our UCE set (Supplemental Information file “BUSCO_UCE_chromosome_Tcas_Psulph”). Results show that within a chromosome, the order of BUSCO genes is not conserved (Fig. 7), with few long segments of synteny within a chromosome. Synteny is greatest between *P. sulphureomaculatus* and the three Polyphaga beetles, and least between Adephaga (*Pogonus*) and *P. sulphureomaculatus*. Interestingly, there is more synteny between *P. sulphureomaculatus* and *Photinus pyralis* (firefly) (Fallon et al. 2018) than between *P. sulphureomaculatus* and *Propylea japonica* (ladybird beetle), the closer lative of the two, indicating that the lineage leading to *Propylea* has undergone many more chromosomal translocation events (Fig. 7 and 8).

**Figure 7.**
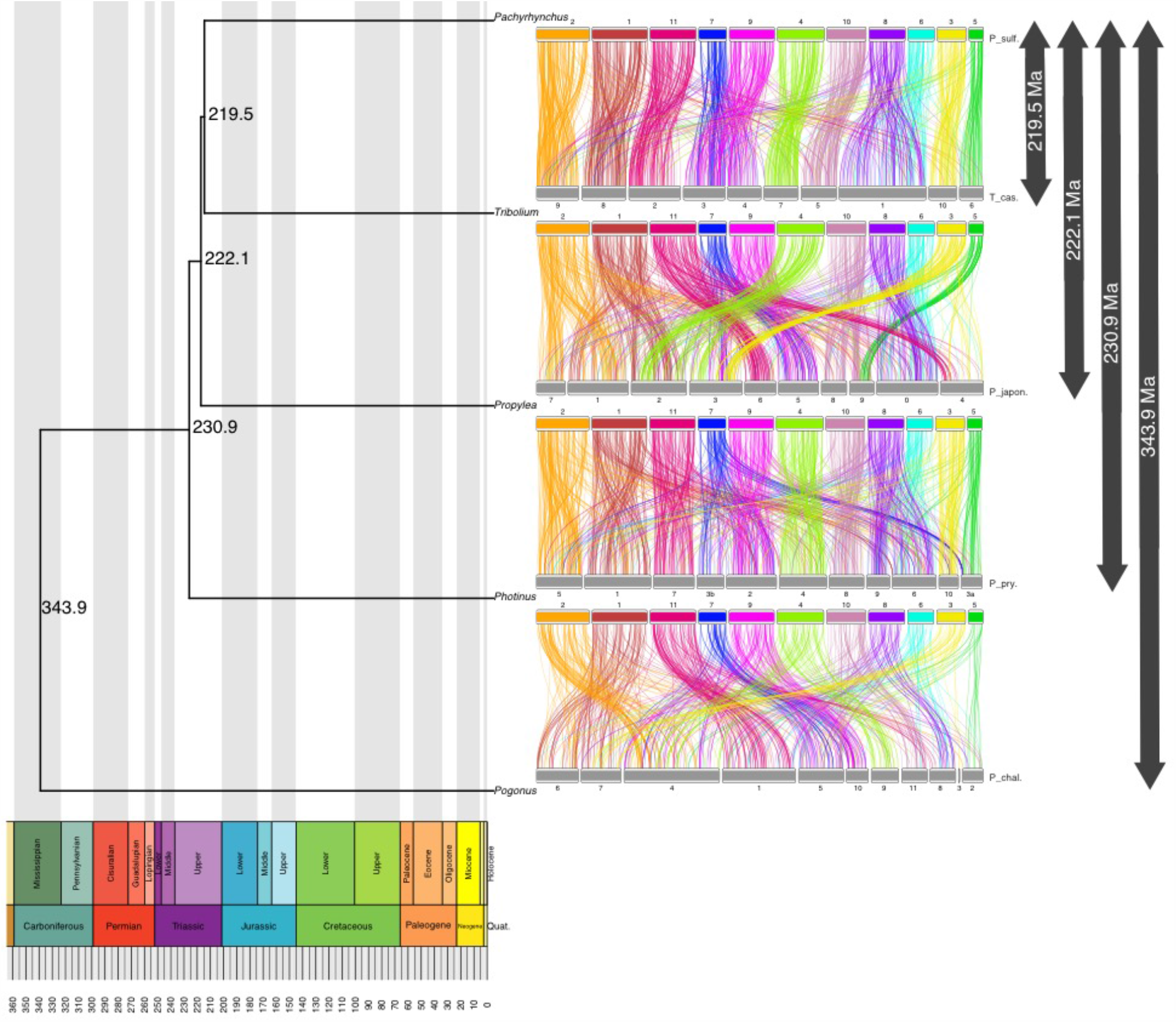
Chronogram and ideograms of the 5 beetle genomes which have chromosome level assemblies.

**Figure 8.**
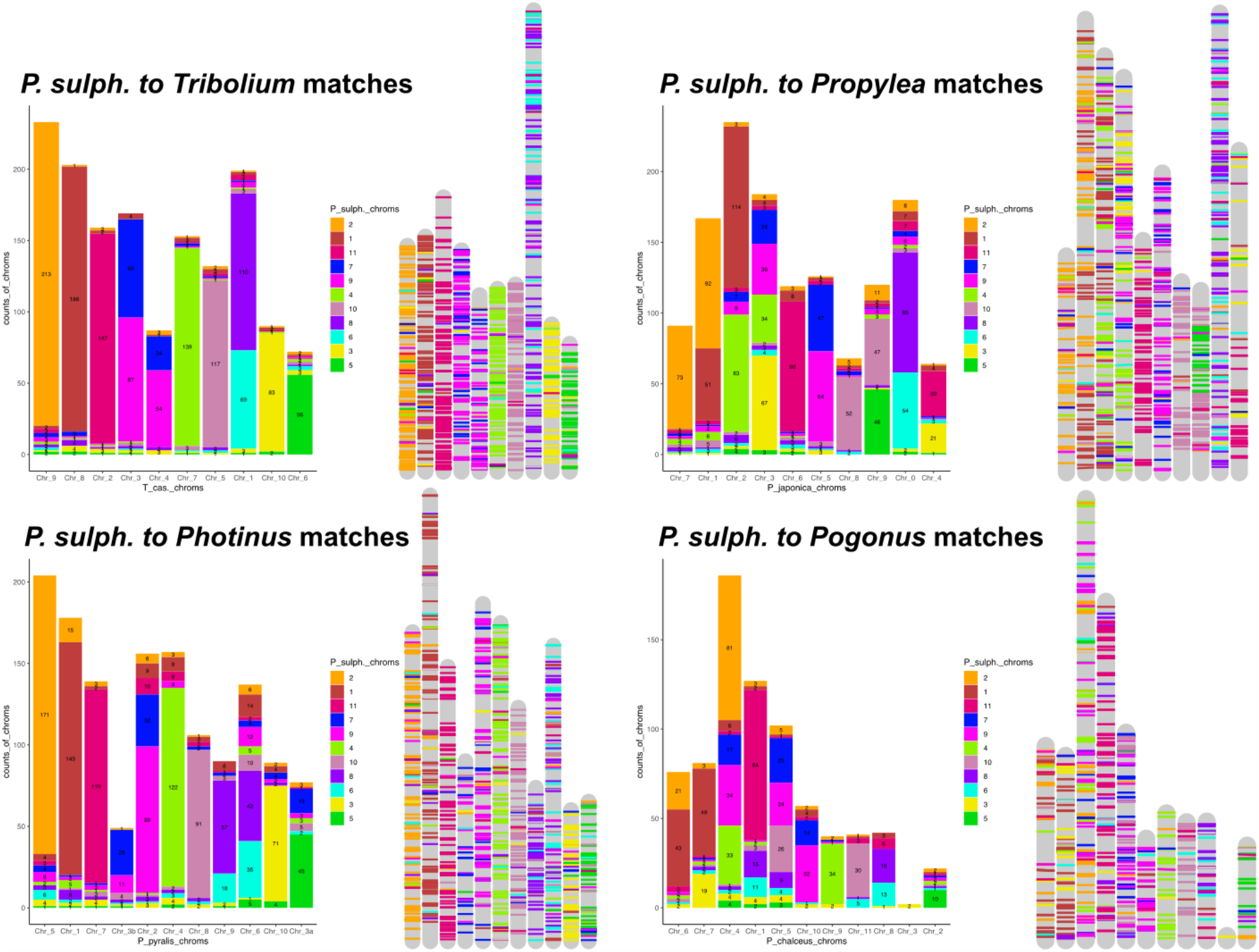
Stacked bar plots and chromosome mappings of BUSCO genes’ placements. Colors correspond to *P. sulphureomaculatus* chromosomes. The numbering scheme of chromosomes matches the names found in the genome’s fasta file.

We computed the Ensemble Gene Order Conservations (GOC) scores (ref) across all pairwise comparisons for our 34 taxa; results are in Supplemental Information file “GOC_results_matrix.txt”. We recovered 1356 BUSCO Genes for the 70% complete matrix, totaling 546,311 amino acids in length. The phylogeny recovered the same clades as in Misof et al. (2014). The GOC scores tended to vary with phylogenetic distance between taxa. For example, the hemipterans *Triatoam rubrofasciata* and *Rhodnius prolixus* (Reduviidae) scored 0.73, and between *R. prolixus* and the pea aphid *Acyrthosiphon pisum*, 0.01. Results from the Mantel test between phylogenetic distance and GOC score found that the two variables were correlated (P≤0.001).

## DISCUSSION

The combination of long-read DNA and Hi-C sequencing was successful in resolving a large and highly repetitive insect genome. To date, this is the largest insect genome and one of the largest arthropod genomes assembled to chromosome scale, the horseshoe crab’s (*Tachypleus tridentatus*) being only slightly larger (2.06 Gb vs 2.05 Gb) (Zhou et al. 2020). This is remarkable because the assembly of relatively large and highly repetitive insect genomes into highly contiguous ones such as this was previously unattainable (Li et al. 2020). Those efforts were hindered by repetitive contents breaking scaffolds or misjoining them (Dudchenko et al. 2017, Hill et al. 2019, Li et al. 2020). The unusually large size of the *Pachyrhynchus* genome is mostly due to the inflated proportion of repetitive content, 76.4% of the genome (Fig. 5). Again, highlighting the need for long sequencing reads to span the repetitive content. Here we used a single individual to create both our Hi-C and PacBio libraries. The main advantage over using multiple individuals is little loss of Hi-C reads mapped to the scaffolds; it also eliminates the need for isogenic lines to be established before sequencing. In our previous attempts to assemble a genome for *Pachyrhynchus*, we were greatly hindered by the loss of mappable reads when using multiple individuals. As long read sequencing improves in its capabilities of using a small amount (5–50 ng) of DNA, capitalizing on this combination of Hi-C and long-read sequencing will make it feasible to assemble chromosome scale genomes from single, very small insect specimens (Kingan et al. 2019, Schneider et al. 2020).

The conserved inter-chromosomal synteny (few chromosome translocations) between the beetle genomes is surprising given the divergence times of the different lineages. For example, we recovered chromosomes that have remained 80–92% intact for more than 200 Ma (Figs. 7–8). By contrast, the order of the BUSCO genes inside of the chromosomes are highly rearranged, such as chromosomes 8 and 6 in *Pachyrhynchus* and chromosome 1 in *Tribolium* (Figs. 7–8). This initial finding prompted us to examine whether similar patterns are observed across other insect orders. A characteristic of Lepidoptera is having a high level of synteny across different families (Hill et al. 2019, Wan et al. 2019). When we examine the ages or relative branch lengths between clades, we find that much of the synteny is correlated with clade age (Fig. 9), as well as from the results of the Mantel test (P≤ 0.001). Although Lepidoptera tends to exhibit more synteny given the phylogenetic distance between taxa, there are a few examples other than *Drosophila* in this same time period to compare against (Fig. 9). Currently, chromosome-level genomes are not available for Trichoptera (caddisflies, the sister lineage to Lepidoptera) or early diverging lineages of Lepidoptera. With the addition of these lineages, we could determine whether the observed pattern of synteny conservation is found only in Lepidopteran crown groups or whether it is more widely dispersed across the entire Lepidopteran lineage. Another order with a somewhat similar level of synteny as in Lepidoptera given their divergence times are Hemiptera (Fig. 9), although there are only 2 comparisons with the same level of divergence (Fig. 9). Taxa with much less synteny given their divergence times are Hymenoptera and Diptera (Fig. 9). The finding that synteny tends to decay with age is not surprising; however, there are some insect orders that are more or less syntenic than expected given their age. For example, in *Drosophila*, there is less synteny between members of this genus (∼40 Ma) than across all of Lepidoptera or the Aphidae that we examined. These results of gene order conservation are consistent with research of *Drosophila* topological associated domains (TADs) that showed synteny break points at approximately every 6th gene between *D. melanogaster, D. virilis* and *D. busckii*, which have a similar level of divergence as the *Drosophila* taxa we examined, about 40 Ma of divergence (Renschler et al. 2019). In addition, the chromosomal rearrangement across *Drosophila* tends to occur at TAD boundaries, not inside the loops (Renschler et al. 2019, Liao et al. 2020). In *Anopheles* mosquitos, the TAD structures seem to be associated with cytological structures as well (Lukyanchikova et al. 2020).

**Figure 9.**
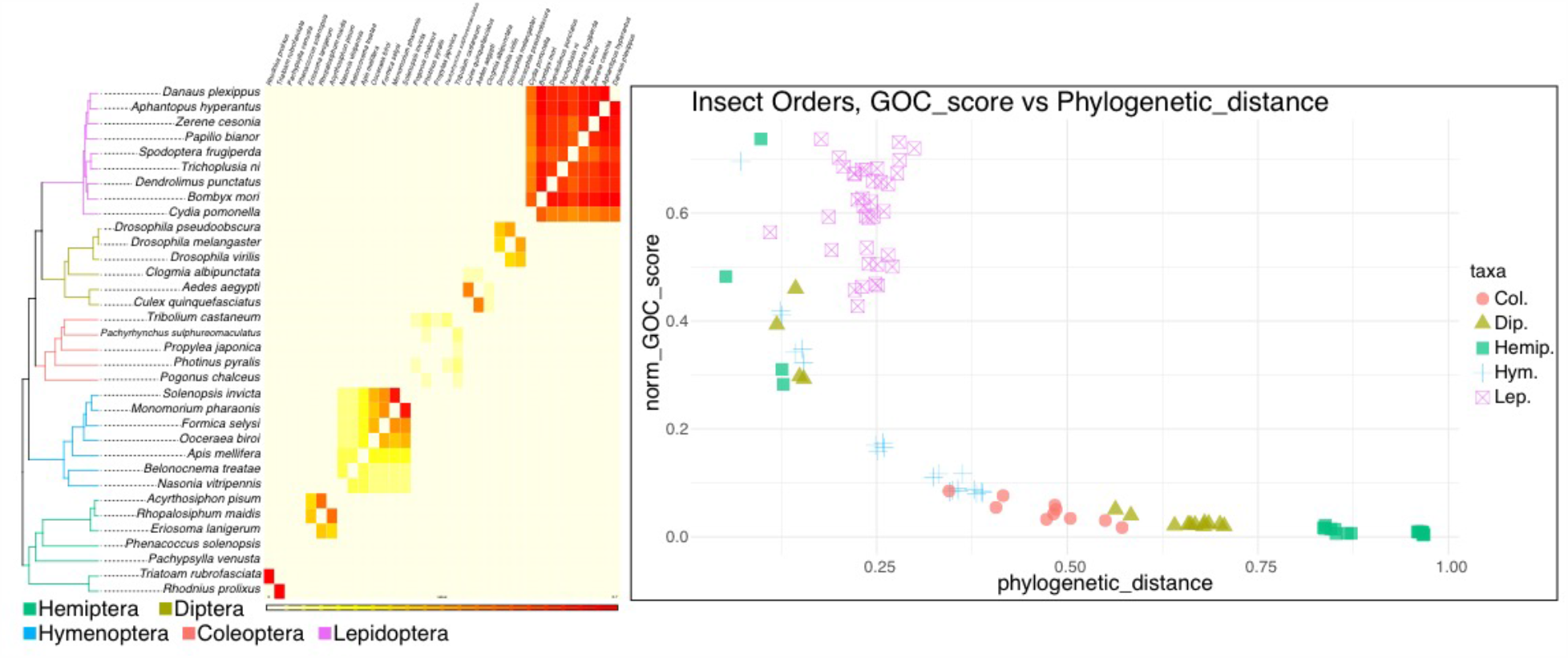
Insecta, gene order conservation score (GOC) plot. Left panel, chronogram, branches colored by order. Heat map of normalized pairwise GOC scores, redder boxes indicate more synteny between pairs. Right panel, within order normalized pairwise GOC score versus phylogenetic distance.

Recent studies in Diptera have demonstrated that syntenic breakpoints tend to occur at the boundaries between TADs (Renschler et al. 2019, Liao et al. 2020, Lukyanchikova et al. 2020). Despite having many breakpoints, with relatively few chromosome translocations, the dipteran chromosomes largely remain intact (Bracewell et al. 2019). In Coleoptera we find a somewhat similar syntenic pattern, in that the chromosomes remain intact while also being highly shuffled (Figs. 7–8). Given the divergence time between our taxa, when translocations do occur, their initial positions are lost due to a high level of reorganization producing a pattern of interwoven segments. For example, chromosomes 8 and 9 in *Pachyrhynchus* and the large chromosome 1 in *Tribolium* (Fig. 8), there are no large syntenic runs of genes or obvious places of translocation. In contrast, chromosome 9 of *Propylea* and chromosome 5 of *Pachyrhynchus* are still largely intact, with the homologous segment of chromosome 5 inserted into roughly the middle of *Propylea*’s chromosome 9. Across the Coleoptera we examined, this was the only fusion event that could be roughly placed. Given the relative amount of reshuffling along other parts of this chromosome, the ability to place the insertion indicates that this was a relatively recent event. This large amount of reshuffling within chromosomes with few inter-chromosomal events contrasts with patterns seen in mammals in which the chromosomes tend to exchange larger blocks of material more readily (Chowdhary et al. 1998, Kemkemer et al. 2009, Deakin 2018, Simison et al. 2020).

Another architectural feature of *Pachyrhynchus*’ genome above the chromosome level includes the Rabl-like configuration of chromosomes, where centromeres and telomeres cluster at opposite/different regions of the cell. These features are important to note because they may serve an important evolutionary function, such as reducing chromosomal entanglements during interphase as well as regulating chromosomal compartmentalization (Mizuguchi et al. 2015, Pouokam et al. 2019). Both major lineages of Diptera, the Nematocera (e.g. mosquitoes and Psychodidae) and Schizophora (e.g. *Drosophila*), have cells with a Rabl-like configuration (Csink and Henikoff 1998, Dudchenko et al. 2017, Matthews et al. 2018). These taxa span much of the phylogenetic distance across the dipteran lineage, and thus this pattern of chromosomal organization may be characteristic of Diptera. We also observe the Rabl-like configuration in *Pachyrhynchus* as well as in the Hi-C map of *Tribolium* (DNAZoo Consortium et al. 2020). Hi-C map observations for the other taxa do not indicate any other obvious cases of the Rabl-like configuration within the Insecta. However, improving the quality of existing Hi-C maps would provide more evidence for this observation because a lack of valid Hi-C reads can obscure this type of chromosomal architecture.

The Rabl-like configuration is not restricted to beetles and flies; it is also found in the yeast genome (Jin et al. 1998, Goto et al. 2001, Mizuguchi et al. 2015, Kim et al. 2017) as well as in wheat, barley and *Brassica* (Santos and Shaw 2004, Mascher et al. 2017, Concia et al. 2020, Wang et al. 2019), and was originally described from salamander cells (Rabl 1885). It is unclear how widespread the Rabl-like configuration is in Coleoptera. The Hi-C maps of the other beetle genomes do not display this formation and are from similar tissue types to what we used (Fallon et al. 2018). It could be that this configuration is only in the Tenebrionoidea and Phytophaga lineages, where it is presently observed. It is assumed that the Rabl-like configuration is found in all life stages, as appears to be the case in Diptera (Dudchenko et al. 2017, Matthews et al. 2018, Lukyanchikova et al. 2020). Additionally, while the Rabl-like configuration is found in different life stages (egg, larva, adult) of *Drosophila*, it is found intermittently in cells, e.g. found in *Drosophila* larvae early G1 and not in late G1 interphase of mitosis (Csink and Henikoff, 1998). Moreover, some organisms’ cells possess a Rabl-like configuration more often during mitosis as compared to other organisms (Idziak et al. 2015). Therefore, what is visualized on the Hi-C map is not that all cells possess the Rabl-like configuration all the time; instead, it is an average of the occurrence in a particular organism’s cells. While the Rabl-like configuration is the predominant chromosomal arrangement observed thus far in Diptera and some Coleoptera, its evolutionary significance remains unclear. Genomic architecture’s influence on diversity, if any, is hindered by the sparse, haphazard sampling of insect genomes. It may be tempting to ascribe patterns to clades (such as Lepidoptera being highly conserved in their genome architecture), but such patterns fade in a broader phylogenetic context or remain to be fully tested. Rather than one to one comparisons, it is more meaningful to describe patterns for a clade in a broader phylogenetic context.

In summation, we have reconstructed one of the largest and most repetitive arthropod genomes. With the combination of Hi-C reads and PacBio long-read sequencing data, we were able to resolve a highly contiguous, chromosome-level genome. We find patterns of genomic architecture, specifically, synteny across Insecta, largely scales with clade age, with some groups, such as Lepidoptera and Diptera, showing subtly different patterns.

## METHODS

### Taxon selection and natural history

*Pachyrhynchus*, from the entirely flightless tribe Pachyrhynchini, is found from the Philippines to Papua New Guinea, Australia, Taiwan, Japan, and Indonesia (Schultze, 1923; Alonso-Zarazaga & Lyal, 1999). They are known for their bright, iridescent and unique elytral markings, which they use as an aposematic signal to warn predators of their unpalatability (Tseng et al., 2014). Members of other weevil groups (e.g. *Polycatus, Eupyrgops, Neopyrgops, Alcidodes*) and long-horned beetles (e.g. *Doliops, Paradoliops*) mimic *Pachyrhynchus*’ aposematic signals to ward off predators. Currently, the Pachyrynchini has 17 known genera, with the majority found exclusively in the Philippines (Schultze, 1923; Yap & Gapud, 2007; Yoshitake, 2013, 2018).

*Pachyrhynchus* Germar, 1824 has the widest geographic range among Pachyrynchini. There are presently 145 species in the genus, of which 93% of are endemic to the Philippines (Rukmane, 2018), with the majority of species having a narrow geographic range, limited to a mountain range, island, or Pleistocene Aggregate Island Complex (PAIC) (Inger 1954, Heaney 1985, Brown and Siler 2014). The general diagnostic characters of *Pachyrhynchus* Germar, 1824 include a head lacking a distinct transverse groove or distinct basal border, entire episternal suture, and antennal scape not reaching the hind eye (Schultze, 1923; Yoshitake, 2012). *P. sulphureomaculatus* Schultze, 1922, is only recorded from Mindanao Island (Schultze, 1922; Cabras et al., 2017; Rukmane, 2018). This species was described from material collected in South Cotabato but has recently been recorded (personal observations of A. Cabras) in other areas of Mindanao (e.g. Marilog, Davao City, Arakan, Cotabato, Mt. Kiamo, Bukidnon). This species belongs to the *P. venustus* group, conspicuous for their large size, prothorax with two dorsolateral spots in the middle a large, oblong spot at the lateral margins, and elytra with oval or oblong spots (Schultze, 1923).

### Collection and extraction of DNA

Specimens were collected near the edge of the road in a secondary forest (HWY 81, Arakan, Cotabato, Philippines [N7.487059, E125.248795]). One individual was used for both in situ Hi-C and high molecular weight DNA libraries. A second individual was used for transcriptome sequencing. Individuals were collected live, then frozen and stored at −80°C until library preparation.

Beetle tissues were dissected carefully to avoid inclusion of contaminants from guts and impurities from chitinous cuticles. Half of the resulting tissues were used for Phenol Chloroform (PCI) based high molecular weight (HMW) DNA extraction for PacBio sequencing (the other half of the material was used as starting material for Hi-C library preparation, see below). Tissues were homogenized on ice using a sterile razor blade. ATL buffer (140 µl) and Proteinase K (60 µl) were then added to the homogenized material and incubated at 65°C for 1 hr. The 200 µl of resulting lysate was used as starting material for the PCI extraction following a PacBio recommended protocol. (https://www.pacb.com/wp-content/uploads/2015/09/SharedProtocol-Extracting-DNA-usinig-Phenol-Chloroform.pdf). Two additional rounds of PCI clean-up were performed to eliminate impurities such as chitin to meet the DNA requirement for PacBio sequencing. In particular, to achieve OD ratios of 1.8–2.0. DNA concentration was determined with the Qubit™ dsDNA HS Assay Kit (Invitrogen corp., Carlsbad, CA), and high molecular weight content was confirmed by running a Femto Pulse (Agilent, Santa Clara, USA).

### In situ Hi-C library preparation

Tissues from the same sample were homogenized using a sterile razor blade on ice. An in situ Hi-C library was prepared as described in Rao et al. (2014) with a few modifications.

Briefly, after the Streptavidin Pull-down step, the biotinylated Hi-C products underwent end repair, ligation and enrichment using the NEBNext® Ultra™II DNA Library Preparation kit (New England Biolabs Inc, Ipswich, MA). Furthermore, titration of the number of PCR cycles was performed as described in Belton et al. (2012).

### Transcriptome library preparation

RNA extraction was performed using tissues from a frozen sample. Tissue was extracted from the prothorax and abdomen with the digestive tract removed. The Monarch Total RNA Miniprep kit (New England Biolabs Inc, Ipswich, MA) was used for extraction. The manufacturer’s protocol for total RNA purification from tissue was followed (cite) (https://www.neb.com/protocols/2017/11/08/total-rna-purification-from-tissues-and-leukocytes-using-the-monarch-total-rna-miniprep-kit-neb-t2010). RNA concentration was determined using the Qubit® RNA HS Assay Kit (Invitrogen corp., Carlsbad, CA), and intact RNA content was confirmed by running a Bioanalyzer High Sensitivity RNA Analysis (Agilent, Santa Clara, USA). The resulting RNA was sent to Novogene Inc. for library preparation and sequencing, from which 12.5 Gbp of data were obtained.

### Genome sequencing and assembly

First, we performed an initial quality control of the in situ Hi-C library using the CPU version of *Juicer* v 1.5.7 (Durand et al. 2016) to determine if enough ligation motifs were present in the sample. To accomplish this, we first cleaned our reads with *fastp* (Chen et al. 2018) to remove sequencing adapters and low quality reads with default settings except for the more sensitive ‘--detect_adapter_for_pe’ setting on. After passing the quality control of having >30% ligation motifs present, we proceeded to sequence the full library at higher coverage. We only considered ligation motifs as this was a de novo assembly without a closely related reference genome to align to the Hi-C reads. The full Hi-C library was sequenced on a paired-end (2×150 bp) lane on an Illumina HiSeq4000. High molecular weight DNA was sent to the QB3 Genomics facility at the University of California Berkeley for sequencing on a Pacific Biosciences Sequel II platform, sequencing one cell with CLR version 2 chemistry (PacBio, Menlo Park, CA, USA).

We used *PacBio Assembly Tool Suite pb-assembly* v 0.0.8 (which includes the FALCON assembly pipeline) to assemble the primary scaffolds. Next, we polished the primary assembly using 3 rounds of mapping the raw fastq reads using *minimap2* (Li 2018) followed by using RACON (Vaser 2017) to help error correct the initial assembly. This was followed by running the *Purge_Haplotigs* (Roach et al. 2018) pipeline to eliminate haplotigs (alternative haplotype contigs) in the assembly. Next, using the CPU version of *Juicer* v 1.5.7, we created a site positions file for the restriction enzyme MboI using *Juicer*’s *generate_site_positions*.*py* script, followed by running *Juicer* until it creates the mapping stats file and a “*merged_nodups*” file. Then we used the 3D-DNA (Dudchenko et al. 2017) pipeline with default settings to correct misjoins and place scaffolds into chromosome groups. After generating a Hi-C heat map, we corrected any assembly errors manually via *Juicebox Assembly Tools* v 1.11.08 (Durand et al. 2016, Dudchenko et al. 2018). After, (Fig 1.) we ran 3D-DNA’s *run-asm-pipeline-post-review*.*sh* to produce a final assembly file and fasta. To polish our final assembly further, we aligned our Hi-C reads to our scaffolds using *bwa mem* followed by *SAMclip* and *SAMtools* ‘view’ (Li et al. 2009) with options ‘-S -b -f 2 -q 1 -F 1536’. After grouping scaffolds into chromosomes, we divided each into a separate fasta (due to memory constraints) and used *Pilon* (v. 1.23) (Walker et al. 2014) in “--fix bases’’ mode as to not break our scaffolds and to fix any homopolymer repeat errors. The resulting assembly was used in all subsequent analyses.

### Removal of mitochondrial/contaminant DNA

To identify scaffolds that contained mitochondrial cytochrome oxidase subunit 1 (COI) DNA, we used BLAT v. 35 (Kent 2002; 2012) using a reference sequence from *Pachyrhynchus smaragdinus* (Supplemental Information file “P79_coI.fasta”) to query our scaffolds. Once identified, these scaffolds were removed. We also used *blast* (Camacho et al. 2008) with the nt database and default settings to identify contaminant (non-arthropod or undetermined) sequences and then removed these from the final assembly. These represented only a handful of sequences.

### Repeat content analyses

To address what is making the genome of *Pachyrhynchus sulphureomaculatus* so large relative to other complete weevil genomes (>85% Benchmarking Universal Single-Copy Orthologs BUSCO Insecta genes), we compared the repeat content of *P. sulphureomaculatus* to 5 other weevil genomes from NCBI (Supplemental Information file “NCBI_numbers_for_Weevils_used_in_repeatmasker”). We used the de novo *RepeatModeler* v. open-1.0.11 (Smit et al. 2015) repeat set combined with all repbase recs to first model for repeat content. Next, we used *RepeatMasker* v. 4.1.0 (Smit et al. 2015) to annotate and soft mask repeat content. For *Listronotus*, we downloaded the results from Harrop et al. (2020), who used comparable methodologies. We also calculated the percentage of repetitive content (bases soft masked) in a 1 Mb sliding window across the chromosomes in *R* using a custom script.

### Genome annotation

We first cleaned our reads with *fastp* and concatenated the unpaired cleaned reads. We performed 3 different initial reconstructions of the transcriptome: 1) *Trinity* v. 2.11.0 (Grabherr et al. 2013; Haas et al. 2014) de novo assembly using default settings, 2) *Trinity* genome guided assembly, where we first aligned our reads with *tophat* v. 2.1.1 (Kim et al. 2013), 3) *rnaSPAdes* (Bushmanova et al. 2019) de novo assembly. Selecting the *rnaSPAdes* assembly, because it had the most single copy *BUSCO V2* Arthropoda genes (Felipe et al. 2015), we mapped our reads to this soft masked assembly using *HISAT2* v. 2.2.0 (Kim et al. 2019), and formatted a bam file using *SAMtools* ‘view -b -f 3 -F 256 -q 10’. Next, we used *BRAKER v. 2*.*1*.*5* (Hoff et al. 2019) to create an annotated gff. This process used the bam file from *HISAT2* and results from a *BUSC*O search as ‘seeding’ genes to make the resulting gff. In addition, we used the *PASA* pipeline (Campbell et al. 2006; Haas 2008) which used our *rnaSPAdes* transcripts aligned to the genome assembly with *BLAT* (Kent 2002) and *gmap* (Wu and Watanabe 2005). Lastly, we used *EVidenceModeler* (Haas et al. 2008) to evaluate our different annotations using the developers’ recommended weights for each assembly type to produce the final gene model gff.

### Synteny across coleopteran and Insecta chromosome-level genomes

To examine the gene synteny between other Coleoptera genomes, we downloaded chromosome-level genomes from NCBI or supplied form the journal or authors website (Supplemental Information file “NCBI_all_taxa_genomes_list”) (Fallon et al. 2018; Zhang et al. 2020; Herndon et al. 2020; Van Belleghem 2018). We also used the unpublished genome assemblies (*Tribolium castaneum* [GCF_000002335.3], *Bombyx mori* [GCA_000151625.1], *Clogmia albipunctata* [clogmia.6], *Culex quinquefasciatus* [CpipJ3], and *Rhodnius prolixus* [Rhodnius_prolixus-3.0.3]), generated by the DNA Zoo Consortium (<u>dnazoo.org</u>). The assemblies were based on the whole genome sequencing data from (Herndon et al., 2020; *Tribolium* Genome Sequencing Consortium 2008, International Silkworm Genome Consortium 2008, Arensburger et al. 2010, Mesquita et al. 2015) as well as Hi-C data generated by the DNA Zoo Consortium and assembled using 3D-DNA (Dudchenko et al., 2017) and Juicebox Assembly Tools (Dudchenko et al., 2018). Next, we identified the BUSCO v.2 loci, (1658 Insecta gene set) and extracted their coordinates for the single and fragmented loci. We then compared the coordinates of *Pachyrhynchus sulphureomaculatus* to the other Coleoptera genomes. Following, we calculated the number of loci found in *P. sulphureomaculatus* chromosomes and those in the other Coleoptera and calculated the percent conserved within a chromosome. To visualize the shared synteny, we plotted the different pairs using the R package *RIdeogram* (Hao et al. 2020). To help visualize the relationship between the different taxa, we generated an ultraconserved elements (UCE) dataset between the taxa using the PHYLUCE pipeline (Faircloth 2012). We used the loci to help reconstruct a concatenated phylogeny in RAxML (Stamatakis 2014) and calculate branch lengths to render the tree ultrametric. We dated the tree using dates (95% highest posterior density interval HPD) from McKenna et al. (2019) using the R package *ape* v.5.4 ‘makeChronosCalib*’* function (Paradis and Schliep 2019) (see Supplemental Information file “Insecta_Claibrations_table” for dates).

Next, we investigated whether the observed synteny was distinctive within Coleoptera relative to other orders of insects, such as Lepidoptera, in which high levels of synteny between taxa have been recorded (Hill et al. 2019, Ahola et al. 2014). We used all insect genomes (with some exceptions) available from NCBI that were marked as “chromosome” level. (See Supporting Information for a complete list.) We tried to sample evenly across insect orders. For example, we excluded the many *Drosophila* genomes as they are all phylogenetically close relatives, and this would cause over-representation (i.e. we want patterns of chromosomal evolution across Diptera, not just *Drosophila*). Instead, we sampled individual species across the phylogenetic breadth of the genus. In addition, we also gathered genomes from the literature. (see Supplemental Information file “NCBI_all_taxa_genomes_list”.) Next, we identified all BUSCO version 5-beta loci that were single copy and calculated the gene order conservation (GOC) score (see https://m.ensembl.org/) using a custom script (Supplemental Information files “1make_scaff_order_busco_tsv.sh”, and “2busco_GOC.sh”). First, we ordered the BUSCO v5-beta genes by scaffold and position and then identified two genes upstream and downstream from a particular gene. Next, to determine if a set of 4 genes are in the same order in our target genome, they receive a score of 1, 0.75, 0.5, 0.25 or 0 based on whether 4, 3, 2, 1 or 0 genes are in the same order, respectively. Missing genes between the two genomes are discarded from comparisons. This process is repeated along the length of the two genomes. We then summed the scores for the four categories 0–100% and added these categories together (e.g. if 8 matched *sets* were found at 25% and 1 at 100%, the total score would be 5). These total scores were normalized by dividing by the minimum number of genes present in the comparisons. We computed the total GOC scores for all pairwise comparisons among the 34 taxa. Next, to consider the effect of the phylogenetic relationships, we reconstructed the relationship among our taxa using the BUSCO gene sets’ amino acids. We used custom scripts to identify a 70% complete matrix and used *mafft* with 1000 iterations and the “localpair” settings to align the sequences. Next, we used *trimAI* (Capella-Gutierrez et al. 2009) with “automated1” settings to remove ambiguously aligned positions. RAxML-ng with the LG+G8+F site rate substitution model was used to reconstruct the phylogeny for our exemplar taxa across Insecta. We calibrated our tree using the same methods for the beetle tree (above), and calibration points can be found in the Supplemental Information file “Insecta_Claibrations_table”, from Misof et al 2014, Obbard et al. 2012 and Mckenna et al. 2019). This calibration was done to help visualize the data as the subsequent Mantel test did not require an ultrametric tree. Lastly, to test if pairwise phylogenetic distance covaries with pairwise synteny values, we conducted a Mantel test.

## Data availability

Assembly available at https://www.dnazoo.org/

Supplemental Information available on Dryad [link pending submission]

Sequencing data from this project are archived on the SRA under accession ####### [embargoed until 2021-11-01 or publication].

## Acknowledgments

We would like to thank the Ruth Tawan-tawan, Ceso II of the Philippines’ Department of Environment and Natural Resources Region XI for help with the Gratuitous and export permits. We would also like to thank the University of Mindanao for the mobility support, and Milton N. Medina and Chrestine Torrejos of U.M. for help collecting specimens. We would like to thank Zane Colaric of B.C.M., for the help loading the QC library runs. We would also like to thank Sarah Crews of C.A.S. for help with the manuscript text. We were funded in part through NSF:DEB award number 1856402 made to MHVD.

